# Better than maximum likelihood estimation of model-based and model-free learning style

**DOI:** 10.1101/296335

**Authors:** Sadjad Yazdani, Abdol-Hossein Vahabie, Babak Nadjar Araabi, Majid Nili Ahmadabadi

## Abstract

Multiple decision making systems work together to shape the final choices in human behavior. Habitual and goal-directed systems are the two most important systems that are studied in the reinforcement learning (RL) literature by model-free and model-based learning methods. Human behavior resembles the weighted combination of these systems and such a combination is modeled by weighted summation of action’s value from the model based and model free systems. Extraction of this weighted parameter, which is important for many applications and computational modeling, has been mostly based on the maximum likelihood or maximum a posteriori methods. We show these methods bring many challenges and their respective extracted values are less reliable especially in the proximity of extremes values. We propose that using a free format learning method (k-nearest neighbor) which uses more information besides the fitted values e.g. global information like stay probability instead of trial by trial information can ameliorate the estimation error. The proposed method is examined by simulation and results show the advantage of the proposed method. In addition, investigation of the human behavior data from previous researchers proved the proposed method to result in more statistically robust results in predicting other behavioral indices such as the number of gaze directions toward each target. In brief, the proposed method increases the reliability of the estimated parameters and enhances the applicability of reinforcement learning paradigms in clinical trials.

## 1 Introduction

Human decision making behavior is believed to be controlled by multiple systems. Habitual and goal-directed systems are responsible for most decisions and learnings during the human’s lifetime [1,2]. While the habitual system reconstructs habits and automatic decisions, the goal-directed system is involved in planning behavior of individuals. Modeling of the habitual and goal-directed systems has been mapped to Model Based (MB) and Model Free (MF) learnings in the literature of reinforcement learning. In MB learning, a model of environment which is the state transition probabilities are assumed to be recruited in mind and the value of each choice in the current state is calculated based on the model of the environment; on the other hand, in MF system the value of each action in each state is learned by trial and error and is updated without considering any explicit model of the environment. In both models, the choices of subjects depend on the estimated value of actions. Investigations on human beings’ behavior have shown that individuals use a combination of MB and MF learnings to guide their behavior during learning tasks[2–6].

Computational hybrid model has proven to be the best descriptor of subjects’ behavior, in which subjects run both MB and MF algorithms in parallel and make choices according to a weighted combination of the action values. Through this model, just one parameter (*w*) determines subject’s preference towards MB and MF algorithms[7]. In many studies, this free parameter is the relative weight of the two algorithms’ values that combines two algorithms’ values and results in the final choice. It is usually assumed constant for each individual subject throughout the task but can vary across subjects[7–10].

The combination weight (*w*) is a free parameter which shows the interplay between two learning strategies, MB and MF, and is important in understanding human behavior. Consideration of changes in this parameter due to pharmacological or cognitive manipulations or neuropsychiatric conditions will provide important insights for clinical research. Over-reliance on habits, for example, could lead to inflexible decision-making in addiction and compulsion[10,11]. Patients with obsessive compulsive disorder (OCD) show a deficit in goal-directed control and an over-reliance on habits[12]. Wit et.al. show that mild Parkinson disease leads to impaired stimulus-response habit formation [13]. Also, Culberth et.al. show that MB behavior is reduced in schizophrenic patients[14].

In a wider view, there is a growing consensus that computational modeling can also be helpful to understand psychiatric disorders. Computational models can break maladaptive types of behavior into distinct cognitive components, while the model parameters associated with the components can be used to understand the latent cognitive sources of deficits[15]. The share of habitual and goal-directed systems are among such components that can be used to assess and diagnose psychiatric disorders[16]. Therefore, estimation of the models’ parameters in general, and the weight of MB learning, in particular, is important for many applications.

Reliable and precise estimation of the model’s parameters is an essential step in extending the use of modeling to clinical applications. However, due to noisy behavior and confounding factors as well as low sample size, reliable estimation of parameters is a challenge especially for extreme values. In many cases, the model fitting results in extreme values for *w*; further analysis shows that the estimated value of *w* by model fitting is less reliable especially when the other parameters of the model are in appropriate range. Simulations show that the low value of learning rate or high value of the Boltzmann machinery temperature ends in the high error of *w* estimation in model fitting.

Here, we propose that using all available information including behavioral measures of subjects besides the fitted values of model fitting can improve the reliability of parameter estimation in the assessment of the *w* parameter. To validate our approach, we used a two-step decision making task, which has been introduced by Daw et.al. previously; It can dissociate MB and MF contributions on human choice behavior [8]. Simulations showed that using some behavioral indices in the estimation of the *w* by the k-nearest neighbor (KNN) method resulted in a lower error in the retrieval of the real weight. Our results improve the applicability of MB and MF task in clinical trials and also in cognitive assessment protocols.

### 1-1 Daw Task

Daw et.al. have developed a two-step task by which MB and MF components of human choice behavior could be dissociated[7]. This task has been used by several other researchers [9,17–25].The Markov Decision Process model for this task is illustrated in Fig 1. The main point in this task lies in the effect of reward in rare events in the decision of next trial differs between MB and MF approaches. In MB approach, a reward in a rare transition (transition with probability of 0.3) is not attributed to the actions of the first level but in MF approach the transition probability is omitted and the reward is assigned to the selected action.

**Fig 1.**
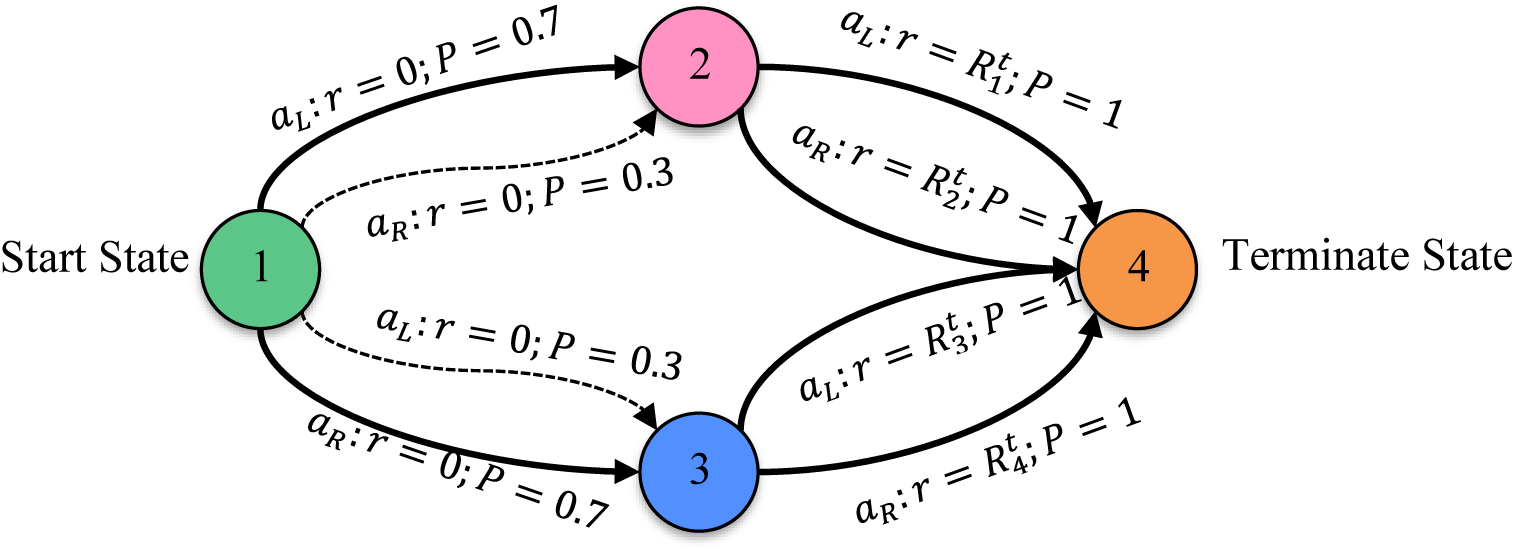
Daw Task MDP model. In all non-terminate states, two different actions are available which are labelled as *a*_*L*_ and *a*_*R*_. Each Start-State action is predominantly associated (with a 70% probability) with one of the second level states. The transitions with 70% probability forenamed “Common”; and those with 30% probability named “Rare”. Any action in state two or three terminate the trial and are associated with different reward probabilities that fluctuate independently across the session by a random walk. In any trial, first action has no reward and the second one results in rewarded or unrewarded trial. Thus, subjects have to make trial-by-trial adjustments in their choice to maximize the probability of achieved reward.

### 1-2 Model Structure

In the modeling of this task, subjects run both MB and MF algorithms in parallel and make choices according to the linear weighted combination of the action values that come from MB and MF systems[7]. This hybrid model has been used by several other researchers[25–27]. Fig 2 shows the flowchart of this model, the parameters of the model and available observations from the human task for each section.

**Fig 2.**
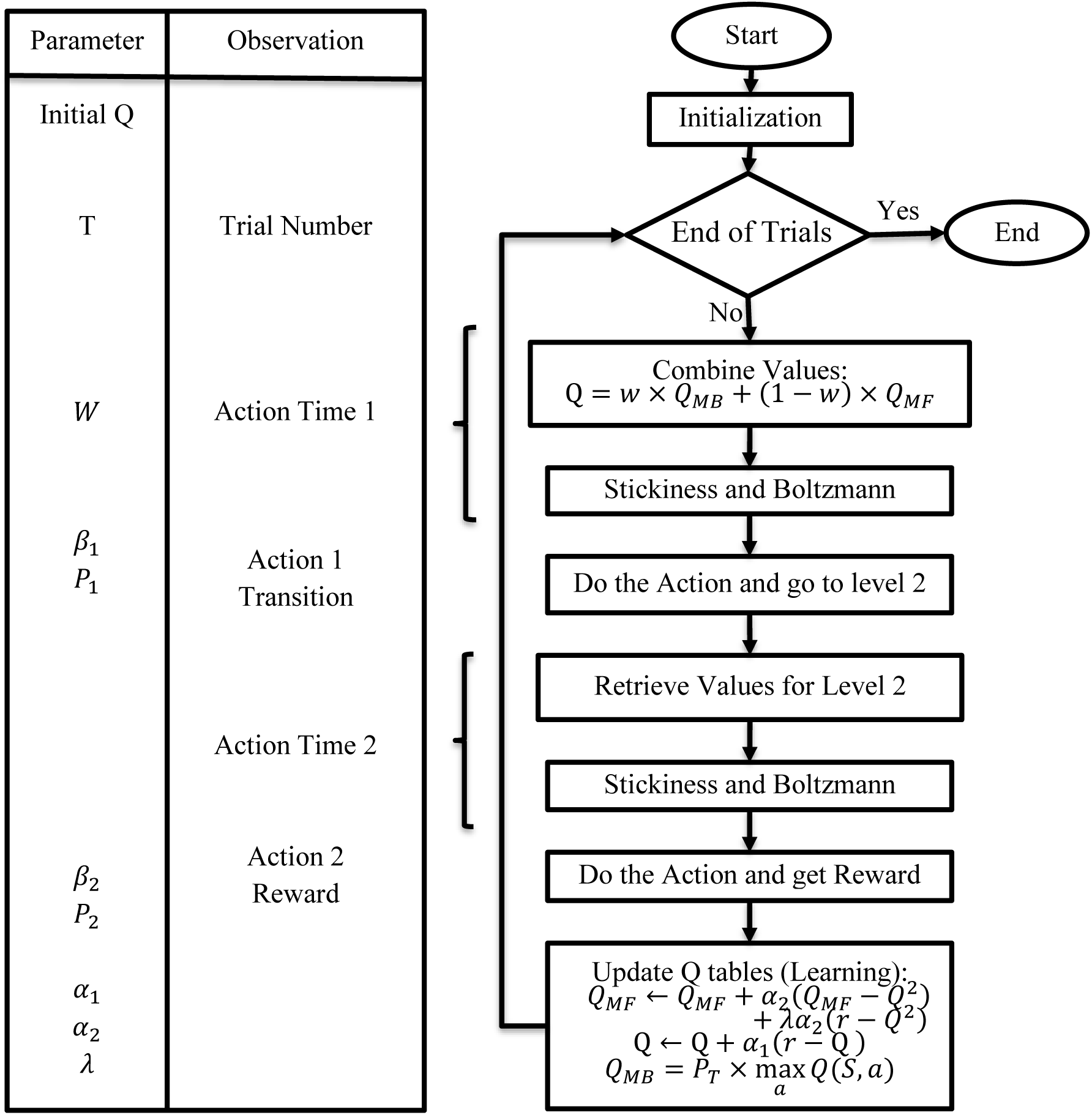
The hybrid model for reinforcement learning. This reinforcement learner is the basic model and we assume that human behavior is based on this model. The parameter box specifies the parameters used in each part of the model. Also, the observation box specifies the available observation from behavior for each part of the model.

In any trial (*t*) the value of each action (a) of first stage was calculated by the weighted sum of MB 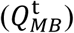 and MF 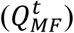 system value (weight: *w*) according to equation (1).

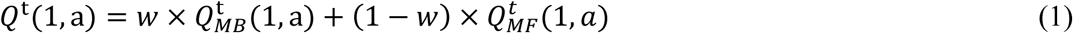

The stickiness to the previous action increases the value of the previous action by adding *P* to the value of previous action (equation (2))

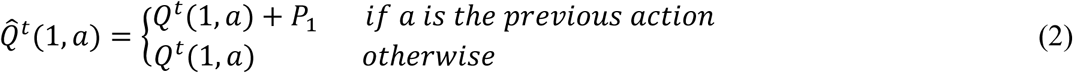

The Boltzmann machine which is a stochastic, biologically-plausible approximation of the maximum operation [28] is used to extract the probability of choosing each action based on their values(equation (3)).

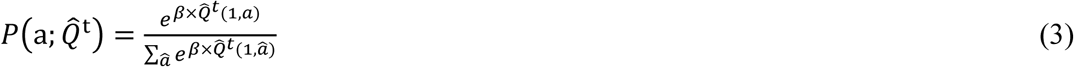

“*β*” is the inverse temperature which controls the trade-off between exploitation and exploration. Due to the non-deterministic environment and its probabilistic nature for rewards, it is usually assumed as a fixed parameter over trials but differs across subjects[7].

In the second stage of the task, corresponding *Q* values in each state determine the probability of chosen action by the same stickiness and Boltzmann machinery. Similar to the approach taken by Daw et. al., the probability of reward for each action of the second stage fluctuates independently across a session according to Gaussian random walks (with the standard deviation of step size: 0.1) and is limited between 0.25 and 0.75.

At the beginning, *Q* values are initialized and the update rules (equation (4)) change the value of each action at the end of each trial. For the second stage of the task, the update rule is the same between MB and MF approaches, but the first state value update is governed by State–action–reward–state–action (SARSA)-λ for MF method and uses the model of environment for update the *Q*_*MB*_. It should be noted that *Q*_*MF*_ and *Q* update rules are applied only to the performed action but *Q*_*MB*_ is updated for both actions.

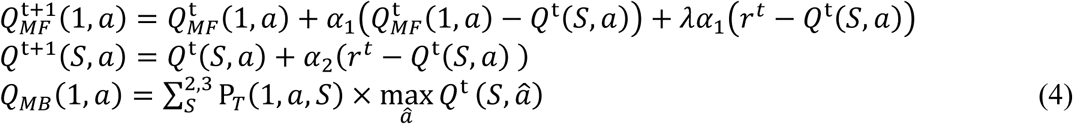

where P_*T*_ is the probability of transition towards the second step state S and calculated by Beta-Binomial Bayesian updating rule according to equation (5) [8].

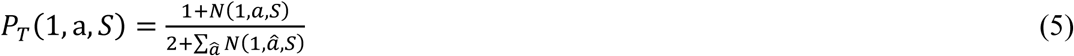

where *N(1,a,S)* is the number of times S state has been achieved by performing the action *a* in the first step of the task.

The initial values for *Q*s are set to zero and the stop criterion is the fixed number of trials, T, which has been set to 150 based on [8] to have faithful comparison condition to human behavior. It should be noted that we also have done all analyses with T equal to 201 based on [7] and the simulation results became clearer. We introduced the model in its most general form above, however alternative models with less free parameters exist in the literature. So, we also used nine versions of this model by fixing some parameters to a fixed value or identical in two stages. These models with different subsets of parameters are shown in Table 1.

**Table 1.** Comparison of model versions

| Parameter Name | combination weight | Learning Rate 1<sup>st</sup> Step | Learning Rate 2<sup>nd</sup> Step | Temperature 1<sup>st</sup> Step | Temperature 2<sup>nd</sup> Step | Eligibility Trace | Stickiness to repeat the same 1<sup>st</sup> action | Stickiness to repeat the same 2<sup>nd</sup> action |
| --- | --- | --- | --- | --- | --- | --- | --- | --- |
| Parameter Symbol<br>Version | $w$ | $\alpha_1$ | $\alpha_2$ | $\beta_1$ | $\beta_2$ | $\lambda$ | $P_1$ | $P_2$ |
| 3ParamV1 | $w$ | $\alpha$ | $\alpha$ | $\beta$ | $\beta$ | 1 | 0 | 0 |
| 3ParamV2 | $w$ | $\alpha$ | $\alpha$ | $\beta$ | $\beta$ | 0 | 0 | 0 |
| 4Param | $w$ | $\alpha$ | $\alpha$ | $\beta$ | $\beta$ | $\lambda$ | 0 | 0 |
| 5ParamV1 | $w$ | $\alpha_1$ | $\alpha_2$ | $\beta_1$ | $\beta_2$ | 1 | 0 | 0 |
| 5ParamV2 | $w$ | $\alpha_1$ | $\alpha_2$ | $\beta_1$ | $\beta_2$ | 0 | 0 | 0 |
| 6Param | $w$ | $\alpha_1$ | $\alpha_2$ | $\beta_1$ | $\beta_2$ | $\lambda$ | 0 | 0 |
| 7ParamV1 | $w$ | $\alpha_1$ | $\alpha_2$ | $\beta_1$ | $\beta_2$ | $\lambda$ | $P$ | $P$ |
| 7ParamV2 | $w$ | $\alpha_1$ | $\alpha_2$ | $\beta_1$ | $\beta_2$ | $\lambda$ | $P$ | 0 |
| 8Param | $w$ | $\alpha_1$ | $\alpha_2$ | $\beta_1$ | $\beta_2$ | $\lambda$ | $P_1$ | $P_2$ |

## 2 Method

The overall structure of our algorithm to extract the *w* parameter is illustrated in Fig 3. When model fitting is used to extract the *w,* the model is initially specified and the set of estimated parameters including *w* is correspondingly determined by maximizing a similarity function between the model’s decision and human decisions. Experimentally, both model structure and objective function affect the performance of model fitting to precisely identify parameter values and this effect becomes clearer when the provided data set is insufficient. This can be due to the limited number of trials, interactions between parameters that make them hard to disentangle or lack of behavior that can be used for the fitting process (e.g., in some Pavlovian conditioning experiments)[29].

**Fig 3.**
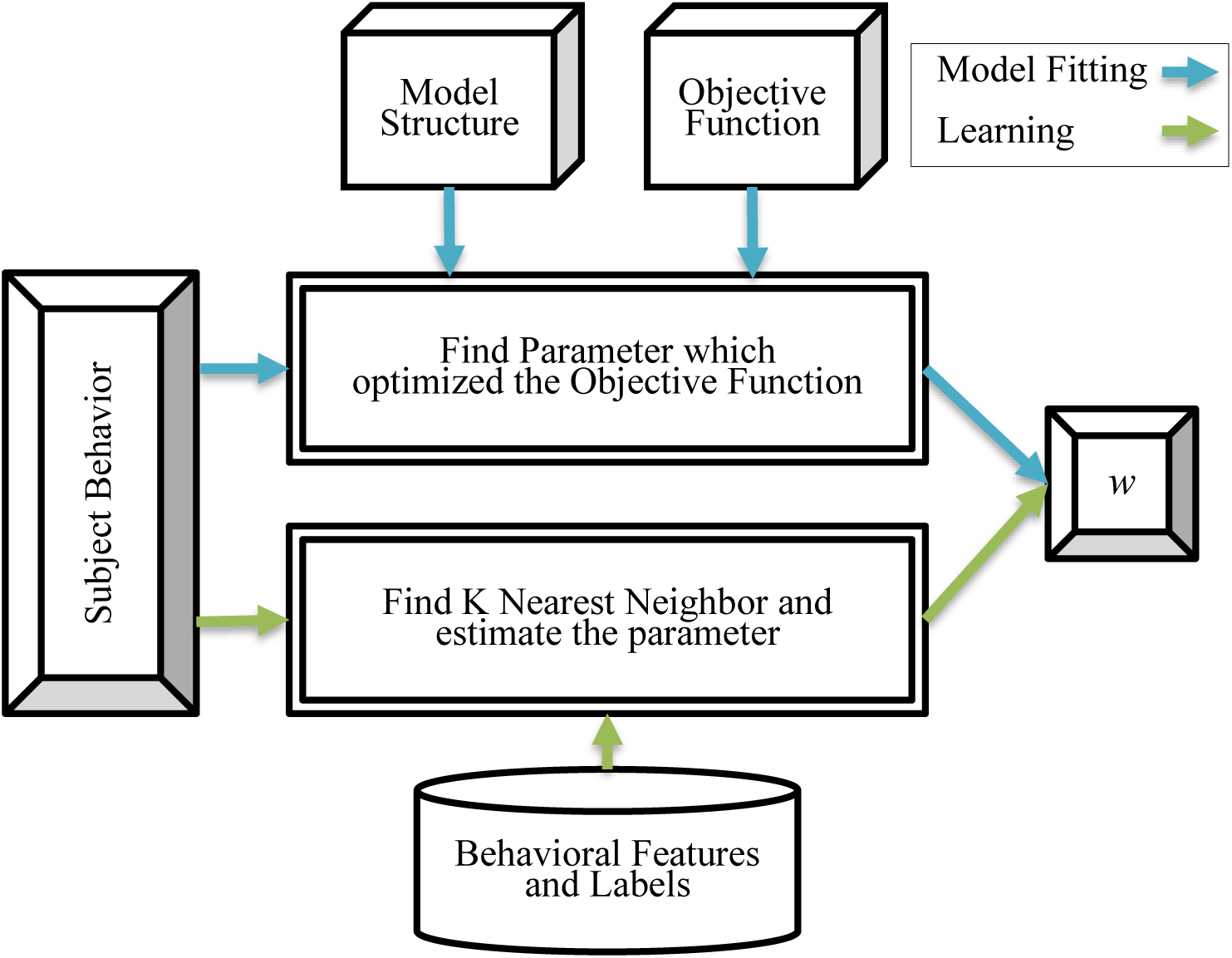
Method for extracting the parameter *w*: Both model fitting and learning get to the observed behavior of subject and the main output is the *w*, the combination weight between MB/MF in the Daw task.

Model fitting is an optimization problem and the objective function is a crucial determinant of the quality of the fitting procedure. Among the objective functions which can be used for fitting the observation to the model, the likelihood function is theoretically the best for fitting the parameters of individual agents [30]. This function tries to maximize the probability of the observed action in the corresponding situation. As a second objective function, to add some prior knowledge, “Maximum A Posteriori” (MAP) method can be recruited which maximizes the probability of a parameter set when an observation is captured [31]. To minimize the objective functions, we used interior-point optimization algorithm with 5 different random start points in the model as mentioned above.

In this paper, we want to use other available information including behavior statistics and indices besides the fitted values of parameter to extract *w* more precisely. In the proposed method, we use a KNN estimator as a learning system to extract the *w* from behavior. KNN uses the feature space of behavior and a dataset of labelled feature vectors which are used to estimate the *w* parameter. The dataset is generated by simulation of the model which has been assumed to be used by subjects.

### 2-1 Proposed KNN Estimator

KNN is a supervised free format learning method and has been used repeatedly by researchers as a good point estimator [32,33]. Different KNN estimators use different distance (neighbor) definitions and different voting methods. We used Euclidean distance in normalized feature space to find *K*-nearest neighbors and then estimations were calculated by the weighted sum of *K*-nearest neighbor values (Equation (6)).

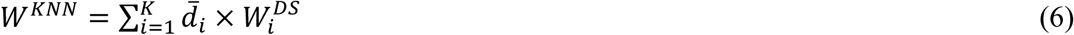

In this equation, 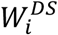 refers to the value of *w* for the *i*^*th*^ neighbor of observed behavior in the database. Also 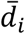 was defined based on the normalized inverse distance according to equation (7).

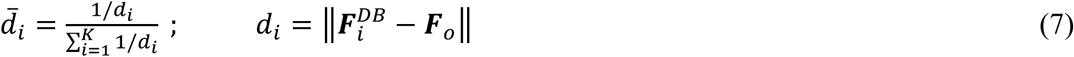

In this equation 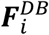 refers to feature vector of the *i*^*th*^ neighbor of observed behavior in the database and ***F***_*o*_ refers to feature vector from observation. Also ║. ║ refers to 2-norm of vector.

### 2-2 Features

We used the following features in the search space of the KNN algorithm: The “stay probabilities” in different situations can be estimated by counting the stays, i.e. same action as previous trial, in the observation data. These different situations are related to the reward value (either Rewarded or Unrewarded) and transition (either Common or Rare) of previous trials which have been commonly used in many papers which used the same task of Daw et. al. Also selection of “Best or Not Best” decision in the first stage of the common trial is another condition for stay probability. The best situation is the one with common transition toward the state with the most probable reward. We used the stay probability in all available situations across three different conditions (listed in Table 2 from 1 to 27).

**Table 2.**
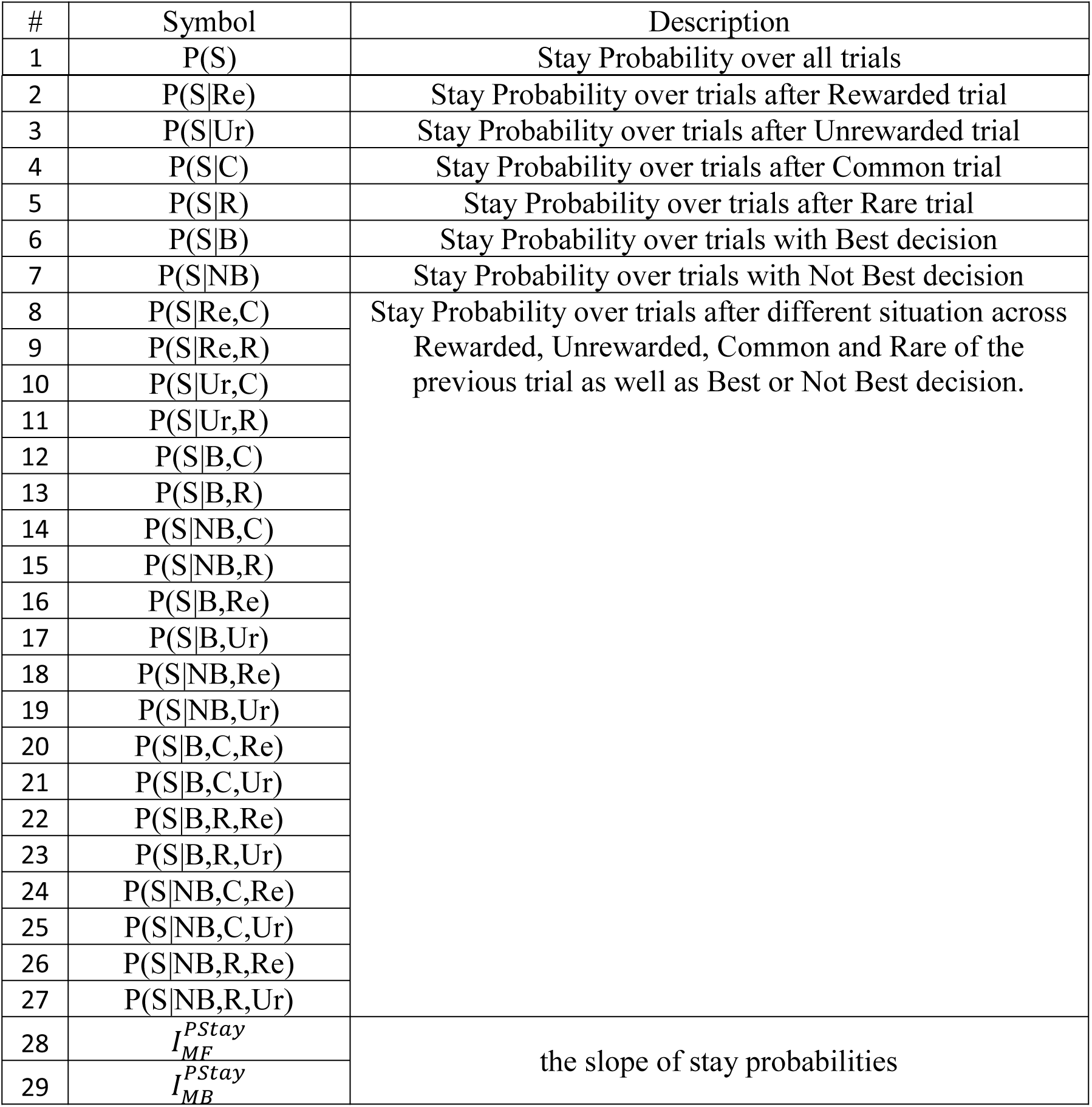
Features extracted statistically based on stay probability

| # | Symbol | Description |
| --- | --- | --- |
| 1 | P(S) | Stay Probability over all trials |
| 2 | $P(S\vert Re)$ | Stay Probability over trials after Rewarded trial |
| 3 | $P(S\vert Ur)$ | Stay Probability over trials after Unrewarded trial |
| 4 | $P(S\vert C)$ | Stay Probability over trials after Common trial |
| 5 | $P(S\vert R)$ | Stay Probability over trials after Rare trial |
| 6 | $P(S\vert B)$ | Stay Probability over trials with Best decision |
| 7 | $P(S\vert NB)$ | Stay Probability over trials with Not Best decision |
| 8 | $P(S\vert Re,C)$ | Stay Probability over trials after different situation across Rewarded, Unrewarded, Common and Rare of the previous trial as well as Best or Not Best decision. |
| 9 | $P(S\vert Re,R)$ | |
| 10 | $P(S\vert Ur,C)$ | |
| 11 | $P(S\vert Ur,R)$ | |
| 12 | $P(S\vert B,C)$ | |
| 13 | $P(S\vert B,R)$ | |
| 14 | $P(S\vert NB,C)$ | |
| 15 | $P(S\vert NB,R)$ | |
| 16 | $P(S\vert B,Re)$ | |
| 17 | $P(S\vert B,Ur)$ | |
| 18 | $P(S\vert NB,Re)$ | |
| 19 | $P(S\vert NB,Ur)$ | |
| 20 | $P(S\vert B,C,Re)$ | |
| 21 | $P(S\vert B,C,Ur)$ | |
| 22 | $P(S\vert B,R,Re)$ | |
| 23 | $P(S\vert B,R,Ur)$ | |
| 24 | $P(S\vert NB,C,Re)$ | |
| 25 | $P(S\vert NB,C,Ur)$ | |
| 26 | $P(S\vert NB,R,Re)$ | |
| 27 | $P(S\vert NB,R,Ur)$ | |
| 28 | $I_{MF}^{Pstay}$ | the slope of stay probabilities |
| 29 | $I_{MB}^{Pstay}$ | |

The stay probability varies between MB and MF system because of the difference in effects of current reward. It has been shown that there are known differences between MB and MF agents in stay probabilities in different situations[7,34]. Furthermore, the slope of stay probabilities, as indices for MF (equation (8)) and MB (equation (9)) behavior [35], were also used as another behavioral indicator in feature space.

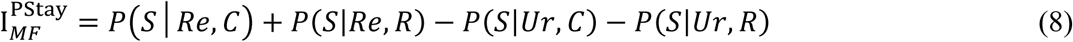

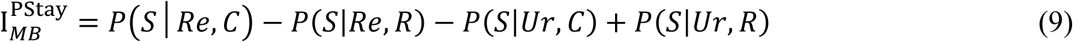

Similar to stay probabilities, these conditions were specified by the outcome of the previous trial.

In addition to the features calculated statistically from behavior, we also add some features from the model fitting procedure. Multiplication of *β* by *w* or (1 − *w*) which are introduced by Miller et.al. as model fitting analysis indices, equation (10) and (11), were added to the feature space [35].

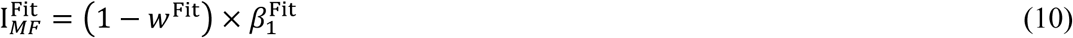

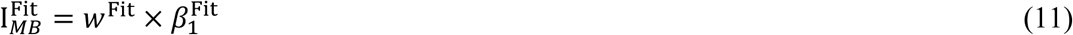

In these equations *w*^Fit^ and 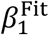 are extracted by best fits according to the Akaike Information Criterion (AIC) by fitting methods which can be Maximum-Likelihood Estimation (MLE) or Maximum a Posteriori (MAP). The estimated MLE and MAP fitting values of some parameters were also included in the feature space. In sum, 10 features extracted by model fitting procedure were added. (Table 3).

**Table 3.** Features by model fitting

| # | Symbol | Description |
| --- | --- | --- |
| 30 | $I_{MF}^{MLE}$ | $I_{MF}^{Fit} = (1 - w^{Fit}) \times \beta_1^{Fit}$ $I_{MB}^{Fit} = w^{Fit} \times \beta_1^{Fit}$ |
| 31 | $I_{MB}^{MLE}$ | |
| 32 | $I_{MF}^{MAP}$ | |
| 33 | $I_{MB}^{MAP}$ | |
| 34 | $W^{MLE}$ | Parameters Extracted by Model Fitting |
| 35 | $\alpha_1^{MLE}$ | |
| 36 | $\beta_1^{MLE}$ | |
| 37 | $W^{MAP}$ | |
| 38 | $\alpha_1^{MAP}$ | |
| 39 | $\beta_1^{MAP}$ | |

### 2-3 Dataset Generated for KNN

As a supervised learning method, KNN needs a training dataset with appropriate labels to function properly. So, we simulated many RL agents with different parameters, with the Daw8Param version of the introduced model, performed Daw’s task in the most complex model and recorded their behavioral observations; simulations were independently run for 80,000 agents and available observations were extracted. Each simulation generated a sequence of trials and related observations which were all labeled by *w*. The random parameters were sampled according to Table 4. Moreover, the 10-fold cross-validation was used for training in hyper-parameter tunings i.e. *K* and feature selection. To remove the estimator bias in extremes, 10000 fully MB agents and 10000 fully MF agents were added to the training dataset.

**Table 4.** Parameters, range and random values for independent agents.

| Parameter Symbol | Parameter Name | Low | High | Random Value |
| --- | --- | --- | --- | --- |
| $w$ | MB/MF weight | 0 | 1 | uniform |
| $\alpha$ | Learning Rate | 0 | 1 | Beta(1.2,1.2) |
| $\beta$ | Inverse Temperature of Soft Max machine | 1 | 10 | $1+9 \times \text{Beta}(1.2,1.2)$ |
| $\lambda$ | Eligibility Trace | 0 | 1 | Beta(1.2,1.2) |
| $P$ | Stickiness to repeat the action | 0 | 0.2 | 0.2×Uniform |

### 2-4 Model the noise in decision making

It has been shown that inclusion of lapse rate for subjects can improve the fitting quality in many psychophysical paradigms. This lapse rate is due to the trials in which subject has not attended and has responded randomly. We included this possibility for agents in simulations. To do so, we simulated agents in a range of noise levels, and at each noise level, a random number of trials, depending on the noise level, were selected for which the decisions of agents were reversed.

## 3 Results

By using random parameters for an RL agent with Daw8Param parameter set, we simulated a wide range of human behavior during Daw’s task. These random parameters were sampled according to Table 4. All fitting methods were applied to observation and AIC was used to choose the best fitted model by each objective function. The model fitting error is calculable for these model fittings because the real value (which is equal to agent *w*) and extracted values are known. While the Mean Absolute Error was used as a point value of error, standard deviation or hinges were used to illustrate the distribution of error. Hinges are the distances between the mean of half data which are below and above the MSE.

### 3-1 Effect of Agent Parameter Set in Model Fitting

Different model assumptions for model fitting can obviously lead to different error levels but what about the agent model version? To investigate this effect, we ran 8000 independent agents with different versions listed in Table 1. For each data set, all model versions and both MLE and MAP model fittings were applied to the observation. The result of these fittings by MLE and MAP is summarized in Fig 4-A and B respectively.

**Fig 4.**
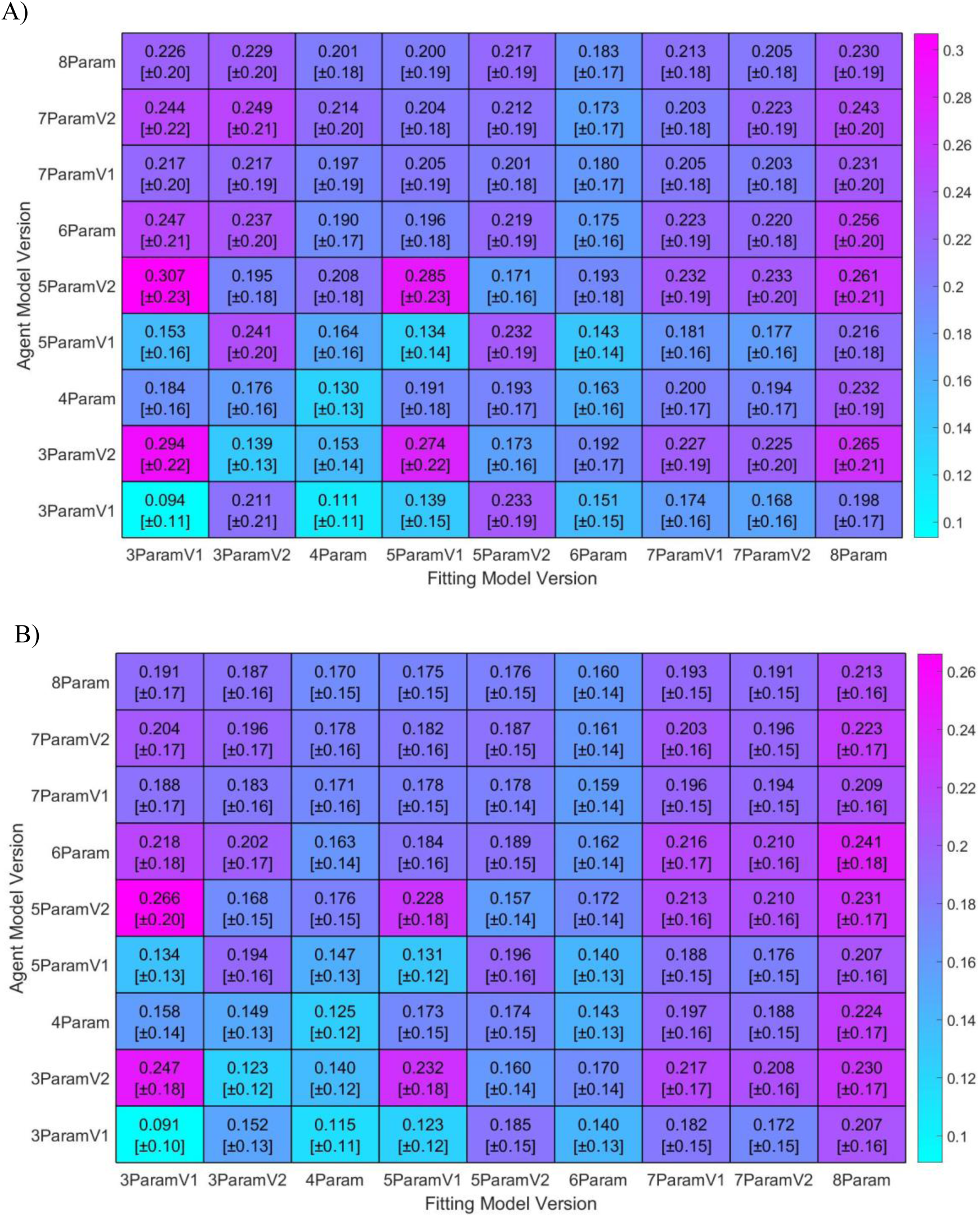
Mean Absolute Error and [STD] of different model version of fitting by A: MLE and B: MAP versus agent model version. Each row is 5000 agents performed the task independently and the column is the result of fitting the model versions to observed behavior.

Based on Fig 4, the largest error was found when the agent was run by 3ParamV1, which has no eligibility trace (*λ* = 0) while the fitting model assumed 5ParamV2 which has a large Eligibility Trace (*λ* = 1). These results were also repeated in 5ParamsV1 version. It can be consequence of that different fixed value assumption for λ make the most errors in models. In fact, *λ* controls the effect of the second stage state-action reward to the first stage action value in SARSA-λ machinery. The behavior of pure SARSA-λ is strongly under the effect of λ value so the information about MB-MF in behavior will be confusing by assuming high λ value for behavior observed from agent whit low λ value and it results in the larger error in the fitting of the w parameter.

Additional to this remarkable point, knowing the model that RL agent has used does not always result in error decrease, especially when the model is more complex. In fact, this happened because of the randomness of behavior. This randomness makes the objective function, multi-modal and increases the number of local minima for more complex model. So, the same model for fitting can be over-fitted for some subjects which result in increasing the error.

Different model comparison methods use information criteria like Akaike Information Criterion(AIC) or Bayesian Information Criterion (BIC) to decide on the quality of model fitting. We selected the best fitted model by AIC and the percentage of selected model is reported in Fig 5-A and B for MLE and MAP respectively.

**Fig 5.**
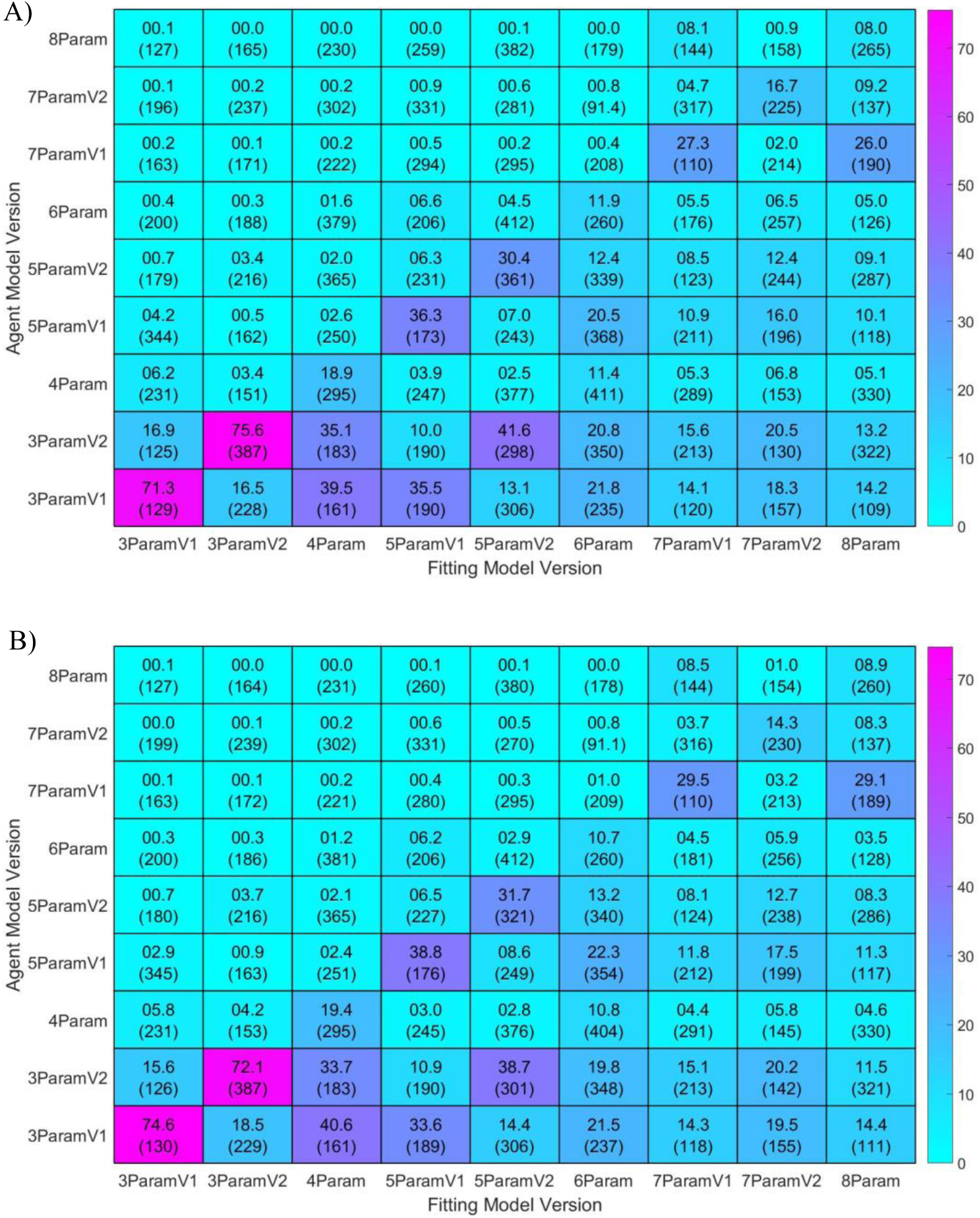
Percentage (Mean of selected AIC) of the best fitting version to different agent versions by A: MLE and B: MAP. The data are same as Fig 4 and each row is 5000 agents perfumed the task independently and the column is the result of fitting the Models for behavior observer.

Most winning models in model comparison are 3Param versions. This happens because of the number of parameters. In sum, using 8Param version for model fitting has no good performance. Because it has less percentage of AIC selection and high MSE due to over-fitting.

### 3-2 Effect of agent learning rate and temperature on model fitting

Assume that the Daw RL agent uses 3ParamV1 version and *w* is sought by best model fitting based on AIC. The question here is whether parameters’ value affect fitting error. To this end, we run 10000 agents with random parameters, according to Table 4, while *α* and *β* are individually assigned with fixed number sets. Fig 6 clarifies the result from this simulation.

**Fig 6.**
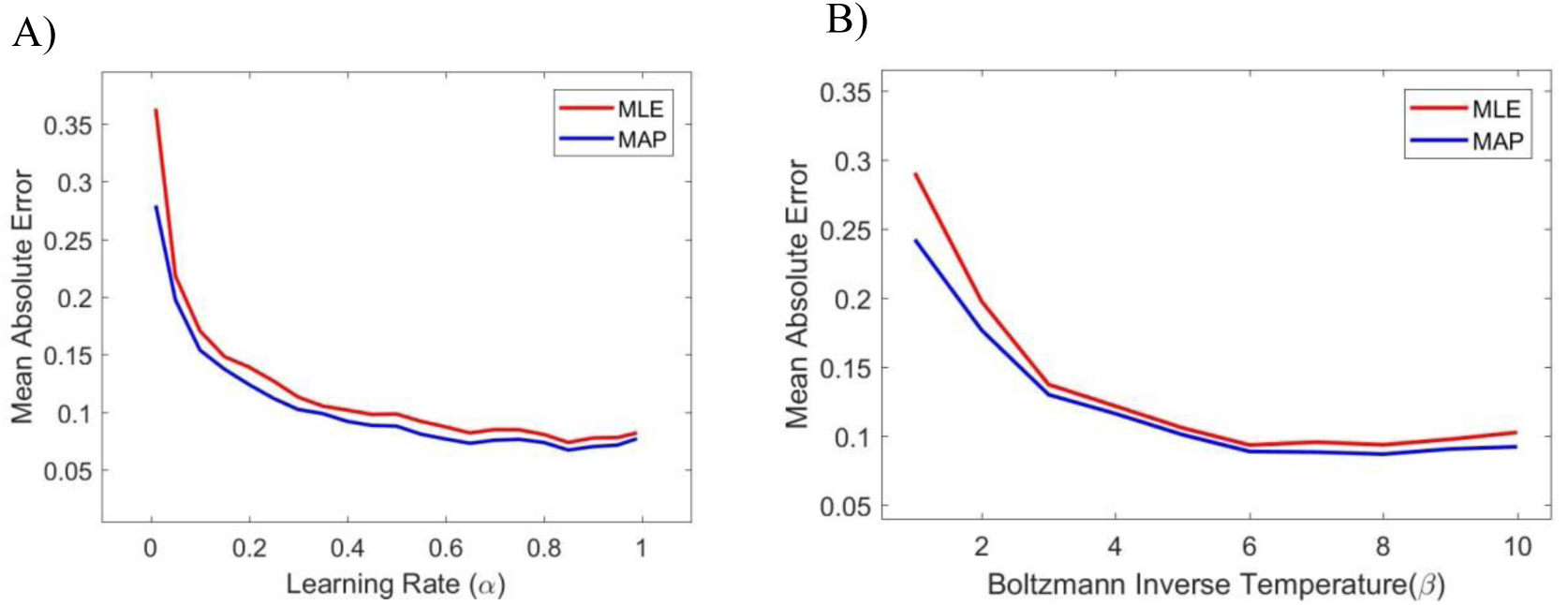
Effect of Agent learning rate (A) and Boltzmann inverse temperature (B) on MSE of fitting. Each point represents 5000 agent that perform the task independently while all other parameters are randomly sampled according to Table 4.

Fig 6-A shows that in low learning rate (α<0.2) the model fitting error is significantly high. In fact, low α means that the agent cannot follow the changes in the environment and this agents’ behavior in both MF and MB systems don’t have a good performance. The behavior is more like random decision making and model fitting faces more challenges and consequently results in greater error. Fig 6-B shows that agents with low Boltzmann inverse temperature (β<3) have high MSE for maximum likelihood fitting method while it decreases for bigger β values.

### 3-3 KNN Parameters

For KNN estimator, the number of neighbors (K) is the only parameter that should be set. In addition to this parameter, the feature selection and feature combination can improve the KNN performance. To achieve the best performance, we found the best K by the exhaustive search to minimize MAE. According to Fig 7.A, the MAE is nearly constant when K is greater than 60 and up to 140 but the value of 101 for K is optimal and was used in all different situations.

**Fig 7.**
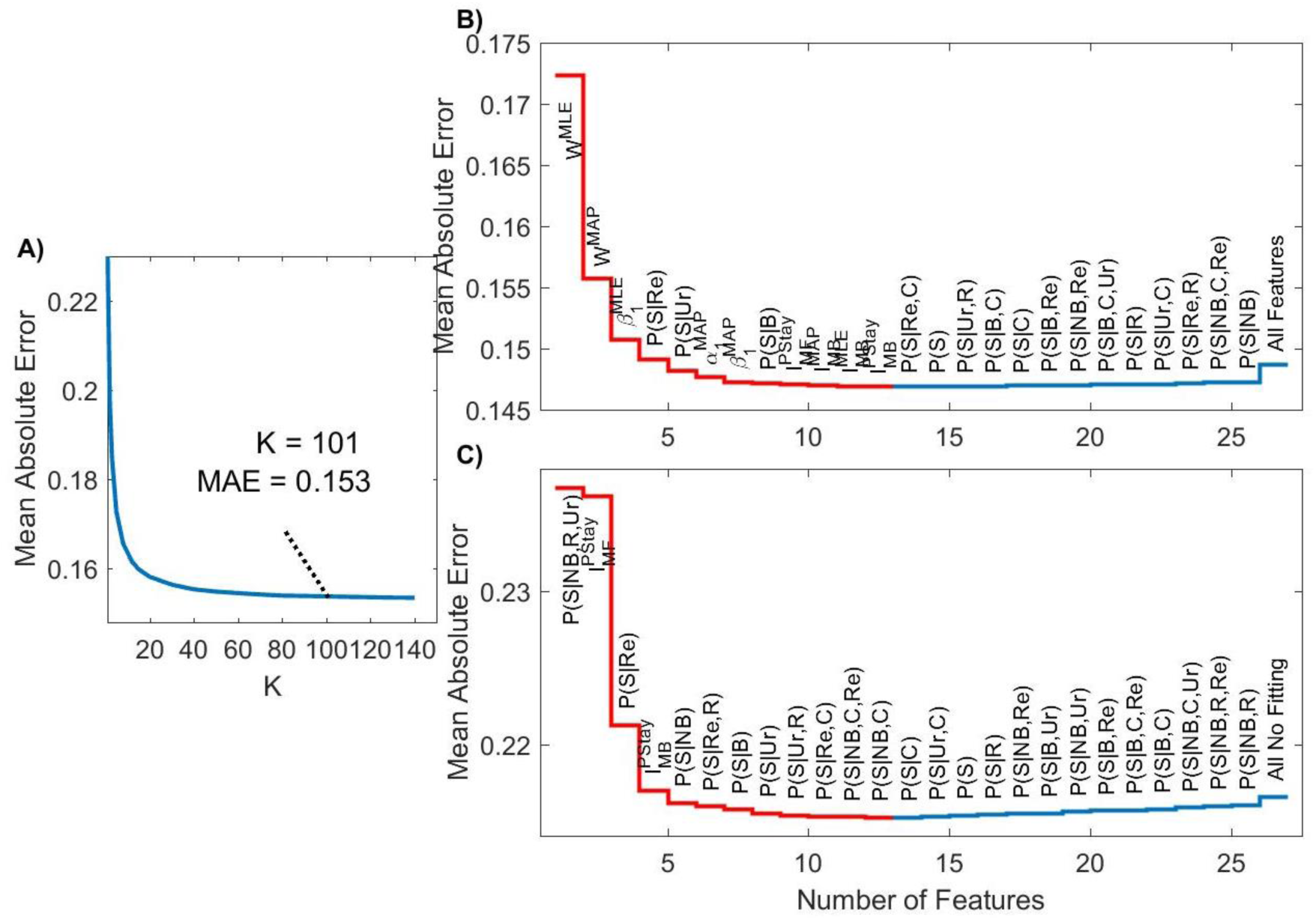
A: Variations of MAE vs K in KNN estimator to find the best K. B and C: Forward selection in different conditions. A: All Features included B: Fitting features excluded. The red steps are selected features for which their inclusion in feature space reduces MAE.

#### 3-3-1 Feature Selection

Selecting good features based on the previously mentioned analyses, can improve the performance of KNN estimator. Forward selection is one of the most commonly used feature selection methods. We used forward selection in two different situations based on the available information and analytics:

1- All features will be computed (needs fitting calculations).

2- Features from model fitting are excluded

Under such conditions for forward feature selection method, different feature subsets can be assumed. Based on Fig 7-B, the subset ℘_sub1_ = {W^MLE^, W^MAP^, β_1_^MLE^, P(S|Re), P(S|Ur), α_1_^MAP^, β_1_^MAP^, P(S|B), I_MF_^PStay^, I_MB_^MAP^, I_MB_^MLE^, I_MB_^PStay^} is selected when all features were assumed to be available. Adding more features just increases the MAE for estimation.

Fig 7-C illustrates the forward selection where total fitting-based features are excluded from features space. The obtained feature subset is then ℘_sub2_ = {P(S|NB,R,Ur), I_MF_^PStay^, P(S|Re), I_MB_^PStay^, P(S|NB), P(S|B), P(S|Ur), P(S|Ur,R), P(S|Re,C), P(S|NB,C,Re), P(S|NB,C)}. Adding any feature results in an increase of the MAE. Although excluding fitting-based features from features space obviously results in higher MSE, it reduces the computational load and the effect of decision noise in results.

### 3-4 Performance

Fig 8 illustrates the scattering of estimated *w* by KNN estimator relative to the corresponding value of agents. The fitted values are the *w* from the best model (based on AIC criterion) by MLE and MAP. The horizontal axis refers to actual *w* which is agent’s *w* and the vertical axis is the output of fitting or KNN estimator.

**Fig 8.**
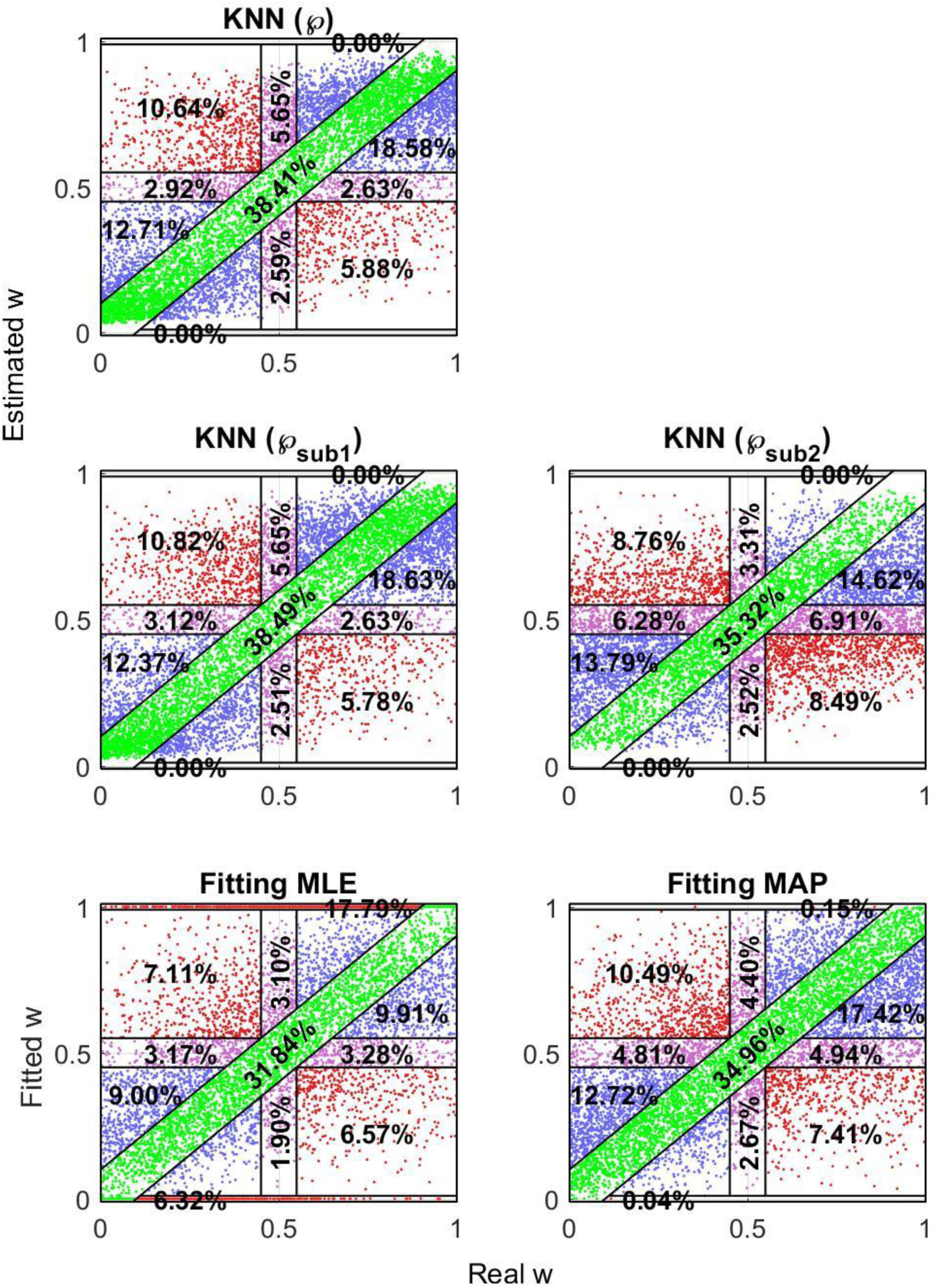
Fitting and Estimation performance. The real W is weighting parameter for 5000 independent RL agents which make the observation by Daw8param model. The fitting extracted w is given by AIC model comparison of model fittings by different methods of fitting also estimated w is the output of KNN in different feature space of ℘, ℘_sub1_ and ℘_sub2_.

To have a deeper view we divide it to different areas. The diagonal area which is scattered points that have a small error (below 0.1). Ideally, this area percentage should be

100. The top and bottom area are those points that reported as common mistake of MLE. We also consider the areas that the dominant strategy changed from MB to MF or reverse, as well as areas that the dominant strategy did not change. In addition, those areas that did not have a dominant strategy has been assumed as the transition areas. The scatter of KNN estimations of w by using all features (℘) and ℘_sub1_ are the nearest values to diagonal. In addition, the scatter illustrates that sticking to the extreme values which is the most problem of fitting methods is solved by KNN.

Fig 9 illustrates error distribution by the normalized histogram of error, the difference between estimated and real values. It is clear that KNN estimation errors are smaller. As mentioned before, the individual difference is an important issue especially in computational psychiatry; and in many cases, percentage of high error is more important than the exact estimation; in the other words, it is important to have an estimation with low error variance. This issue is addressed by bias-variance tradeoff in the literature. Based on Fig 9 the KNN method improves both bias and variance of error against the fitting method. For KNN method, the tail of the distribution is shorter and includes lower values at the tail. It means that variance of error is low and the calculated value of standard deviation (STD) confirms this (Table 5). On the other hand, for KNN methods probability of small error (errors between −0.05 and 0.05) are higher than just fitting methods. In addition, MAE which is reported in Table 5 confirms that the bias of KNN estimation is improved with regards to pure fitting method. Extreme errors in both ML and MAP are relatively higher than KNN based methods. Since it is possible to have extreme values for *w* in clinical conditions, these regions are more important. KNN methods correct these errors and make the method more robust for applications in clinical trials.

**Table 5.**
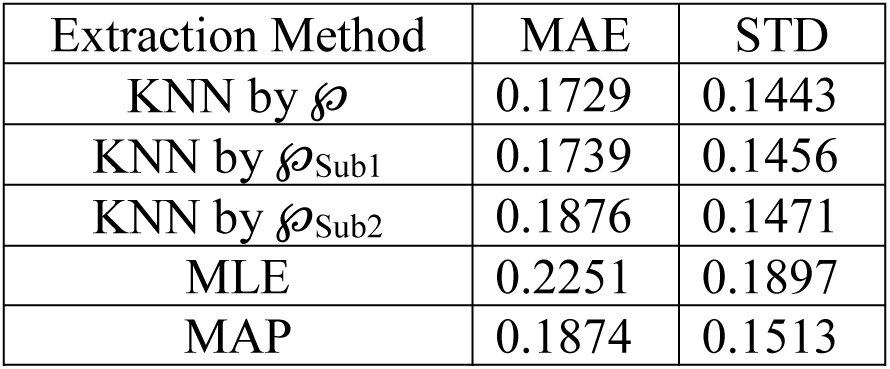
Mean Absolute Error and error STD for w extraction by Model fitting and KNN

**Fig 9.**
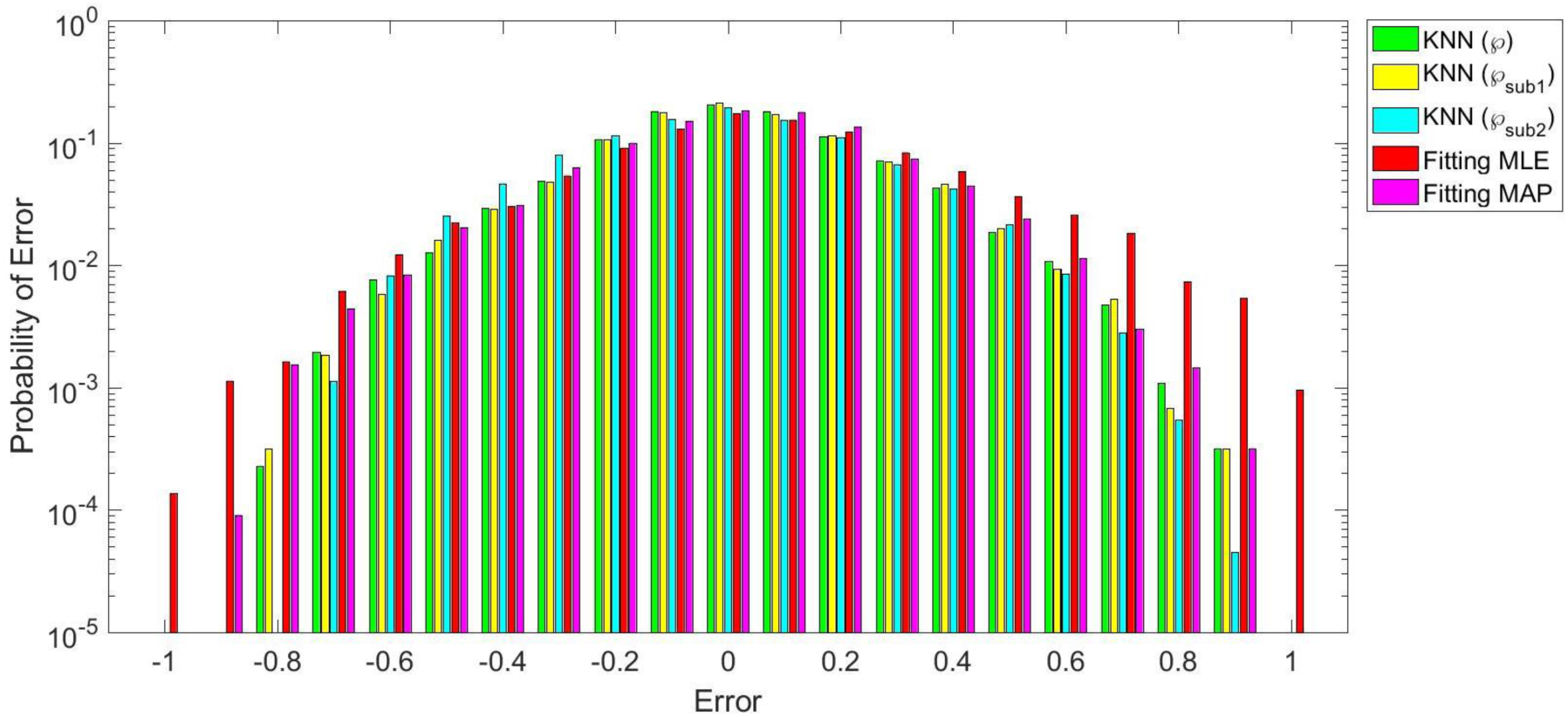
Distribution of error by different models for w extraction. The error which is the extracted value minus the real value, is calculated for 25000 independent agents that makes the observation by Daw8param model version. The extracted w is given by AIC model comparison of the model version by different method of fitting also the output of KNN calculated in different feature space of ℘, ℘_sub1_ and ℘_sub2_.

### 3-5 Noise in decision making

To investigate the effect of human mistake or lapse rate in doing actions, we run the simulation with different probability of additive noise in decision making. For each value of noise ratio, 10000 independent RL agents perform the task and both MLE and MAP fitting were applied on the observation by different models. AIC was then used to select the best fitted model for each individual agent’s behavior. Fig 10 illustrates the difference of fitting methods against KNN estimation method. The KNN estimation is applied in two different conditions listed in feature selection section.

**Fig 10.**
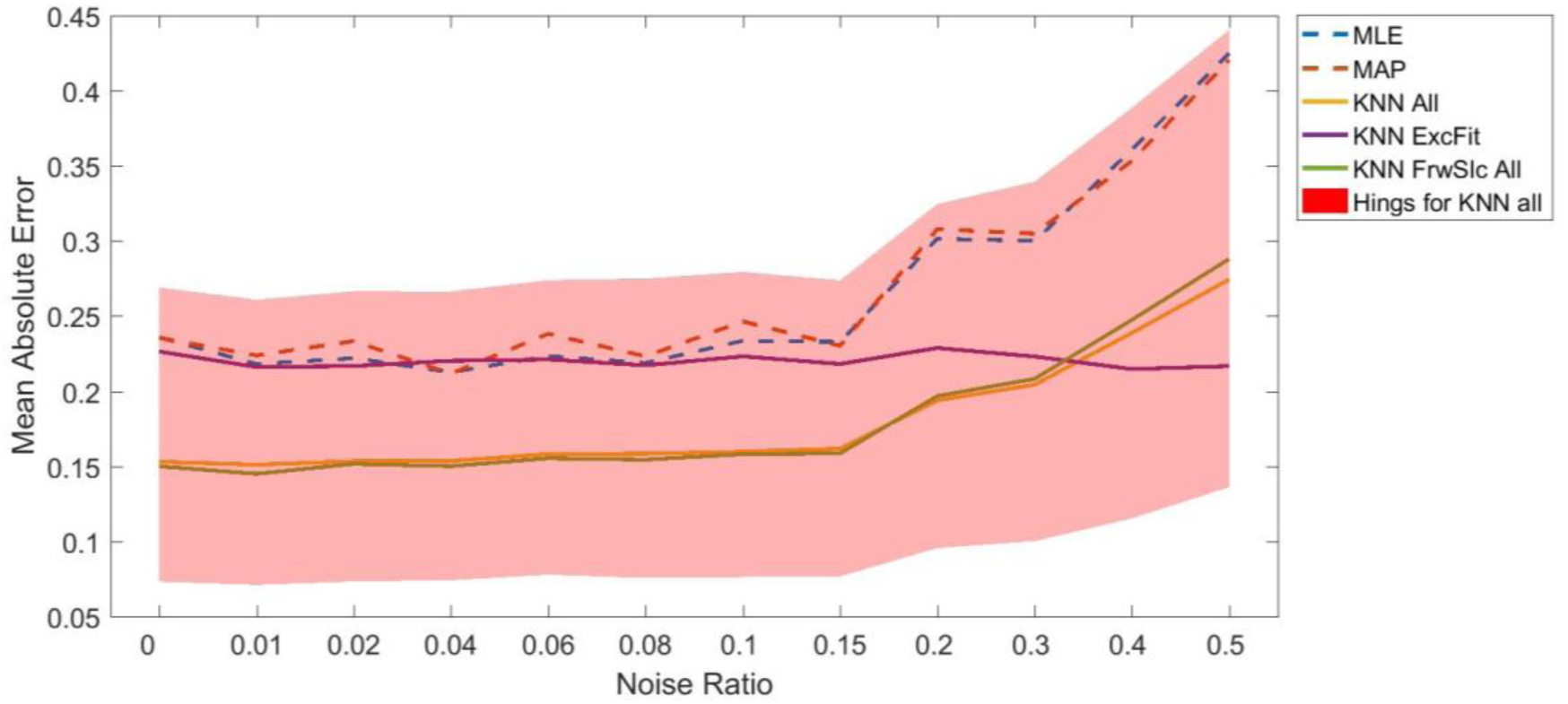
MSE of Extracted w by KNN and Fitting in presence of noise

Based on Fig 10, it is clear that the KNN methods are more robust than fitting methods towards these random decisions, especially when features used in fitting data are excluded from feature space. In fact, the use of statistically extracted features in KNN method, make it more robust to the noise. In the fitting method, however, each decision in trial *t* influences all subsequent trials due to reinforcement learning model.

## 4 Experimental data analyses

The two-step task of Daw et. al has been previously used to investigate the correlation between gaze data and the *w*. [8], where 5Param version of model has been used to extract the w value by MLE. KNN estimation was used to extract the combination weight from their data in the current investigation. Fig 11 illustrates the difference between the “Fitted *W*”, used by Konovalov & Krajbich [8] and the “Estimated *W*” which are the result of our estimation method (KNN by ℘_sub1_ feature space).

**Fig 11.**
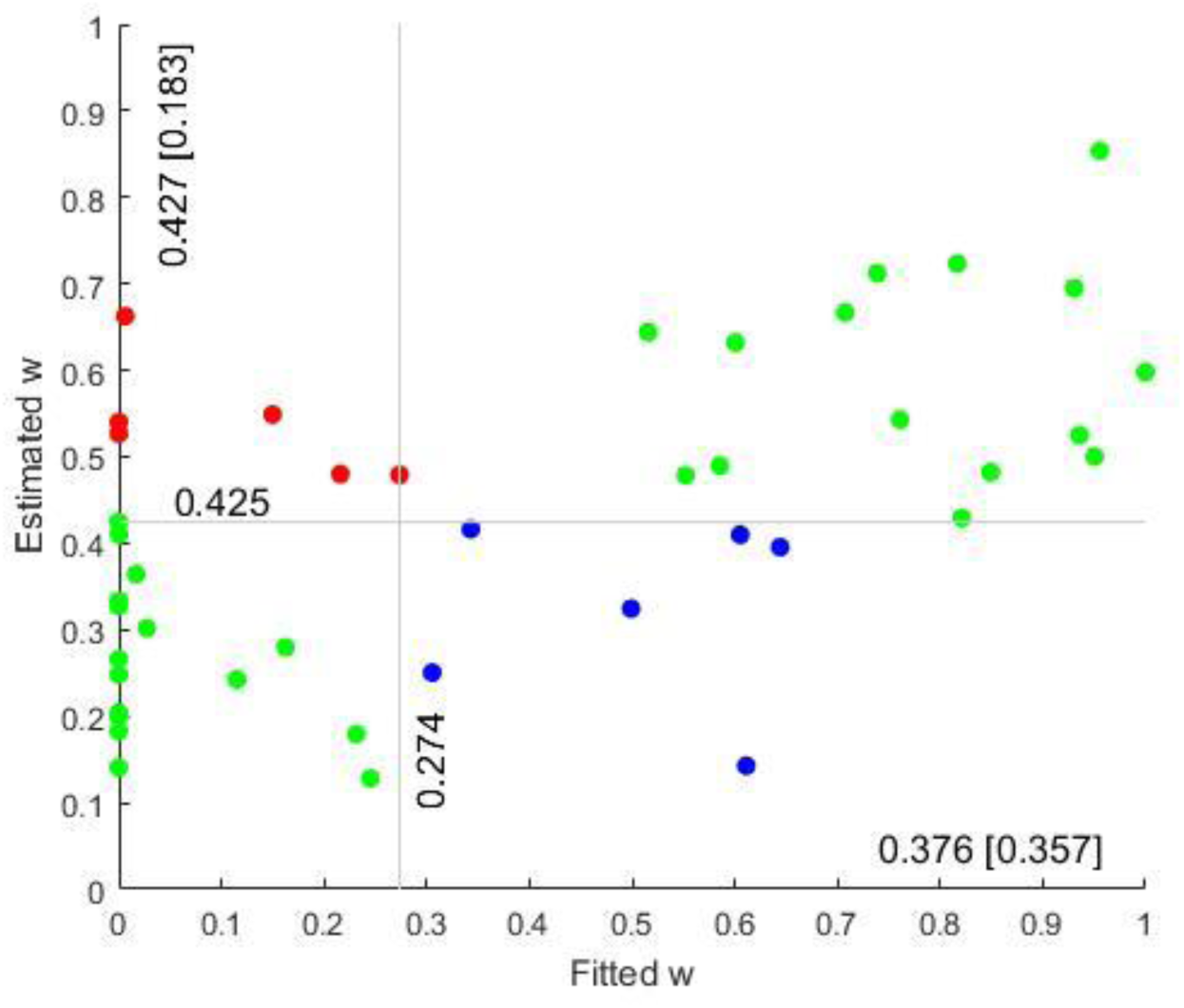
Estimated w vs Fitted w. the green subjects are in the same group by fitting and estimation but the red subjects are MF by Fitting and MB by Estimation also the blue subjects labeled as MB by Fitting and MF by Estimation. (w=0 for pure MF and w=1 for pure MB)

To illustrate the differences between MB and MF behavior, Konovalov et.al. split subjects into two groups based on the median *w* (0.3). In all analyses, ‘model-free’ and ‘model-based’ labels were used for the two groups defined by this median split.

This grouping is first checked by mean value of P-Stay in groups as a behavioral data. According to Fig 11, groups changed for some subjects when the estimated *w* was used instead of fitted *w*. The first question arising then is which groups are more consistent with behavioral observation? To answer this question, the stay probability was extracted through different groupings as well as with group of subjects labeled differently between fitting and estimation (Fig 12).

**Fig 12.**
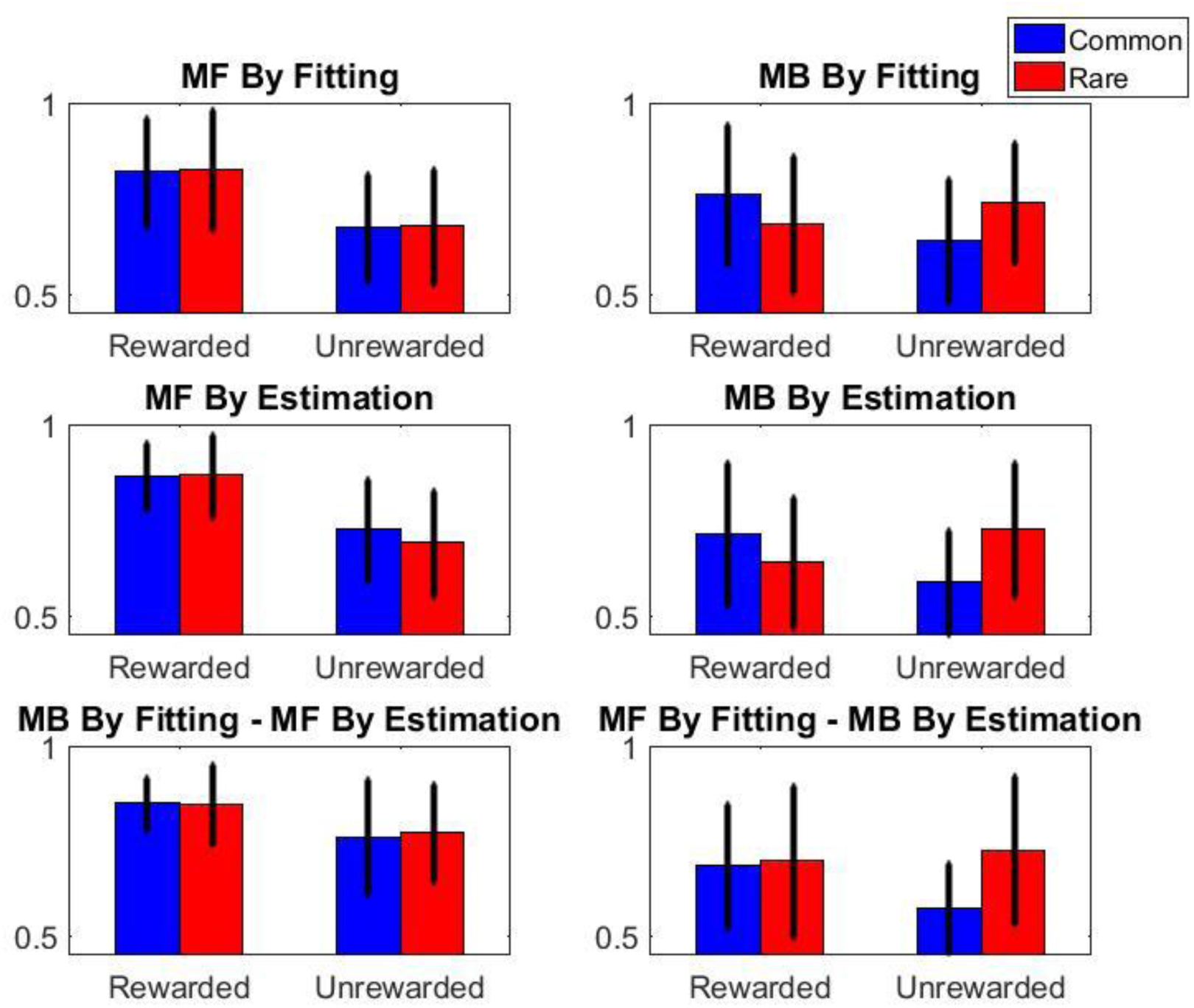
Stay probability for MF and MB groups. The probability is calculated for each subject and the mean of the calculated value for each group and STD is plotted.

Based on Fig 12, both fitting and estimated groupings are consistent with prior findings. However, while label assigned by Estimation for groups that are differently labeled is consistent with prior findings on stay probabilities, fitted values show discrepancies. For six subjects that were labeled as MB by fitting and as MF by estimation (blue subjects in Fig 11), the stay probability in trials after rewarded and unrewarded trials is the same in different transition situations (Common or Rare transition in previous trials); This behavior can be attributed to neglecting the transition (which is the main specification of MF subjects). This subject will therefore be a better candidate for MF rather than MB label and consequently the *estimated w* is better than that obtained by just fitting.

All the analysis given in the first part of the report by Konovalov & Krajbich [8] was checked by new group labels. While the main analysis results did not change, significant level improvements were observed. An outstanding result of gaze data analyses is the insight into difference between gaze number distribution of model-based and model-free groups [8].

Based on Fig 13, the grouping by estimated *w* make the difference between MB and MF groups clearer. Model-based subjects were also more likely to look at only one of the symbols before making their first-stage choice (The average for one gaze is 54 vs. 43 by Fitted *w* grouping and 55 vs. 41 by Estimated *w* grouping). MB and MF groups had different distributions for the number of gazes per trial. Statistical test by both grouping demonstrate the significance of the difference (the P value is 3.24e-13 by fitted *w* grouping and 4.08e-13 by estimated *w* grouping; Kolmogorov-Smirnov test done over all observations). Moreover, for subjects assumed MF by Fitting but labeled as MB by Estimation, the distribution is closer to MB group. However, for those subjects that assumed MB by fitting but labeled as MF by estimation, the distribution is again closer to MB group.

**Fig 13.**
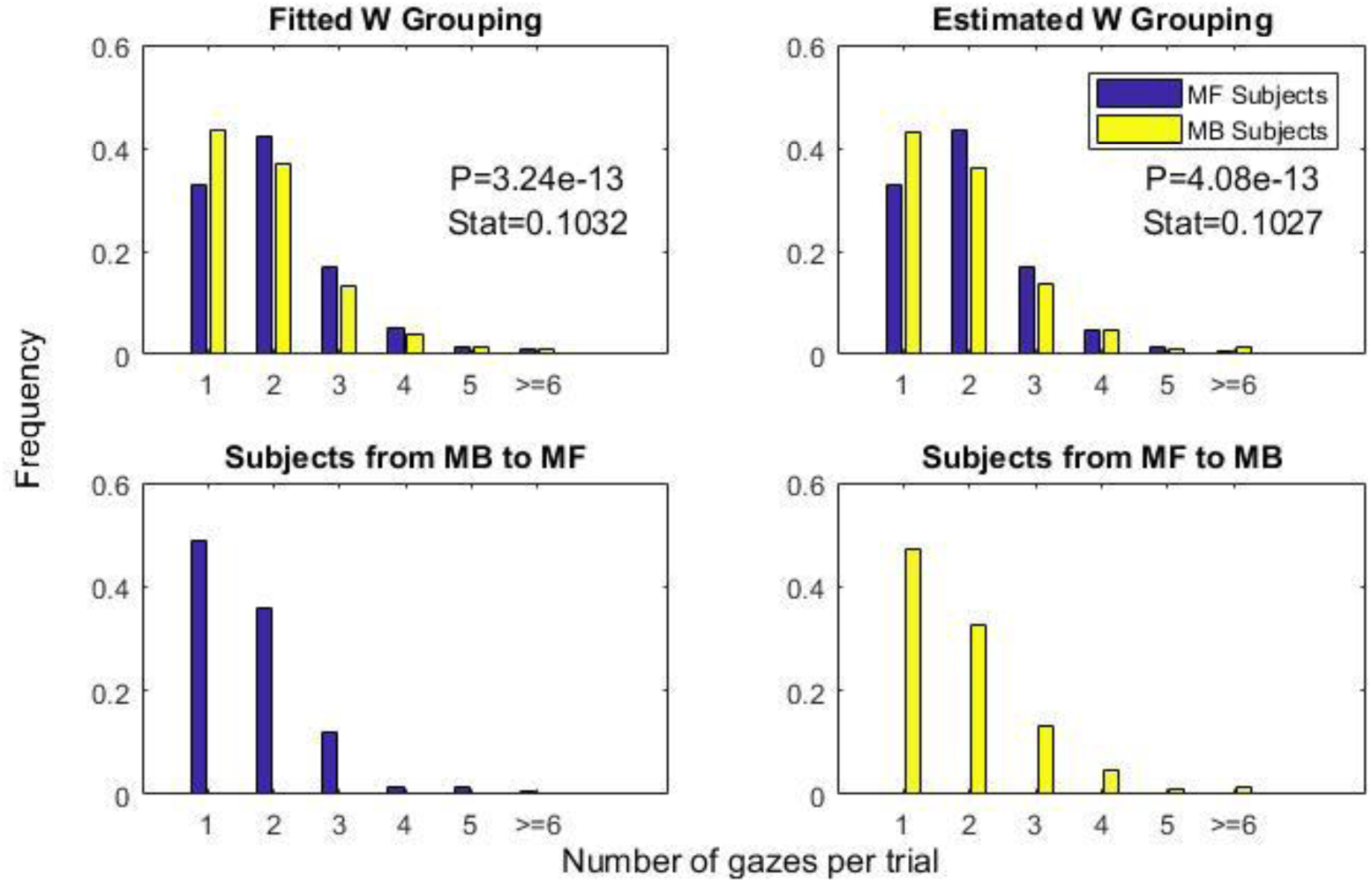
Empirical distribution of number of gazes per trial for different groups. The P value and statistics for the two-sample Kolmogorov-Smirnov test of difference distribution are reported.

## 5 Discussion

Assessment of habitual and goal-directed behavior using reinforcement learning models is a necessary step in translation of the task of Daw et. al. to the clinical trials. We used nine different versions of the main model which are nested models of the most complex used model. Our analyses specify that complex models over-fit to the observation of standard Daw task, especially due to randomness in agent’s decision making. Simple models with wrong assumptions, on the other hand, result in greater errors. Moreover, when prior knowledge was not assumed for the fitted parameters (mainly *w*), the fitted values stick to the extremes of the parameter range. Having a limited number of trials also worsens the problem. Such problems in model fitting make the fitted parameters unreliable.

In this paper, we proposed to use the behavioral information by KNN to estimate a parameter instead of just fitting the model. To have a performance measure, the weighted combination of MB/MF learning was simulated in an agent performing the task and the weight parameter was then estimated to obtain the estimation error. The best performance, which is mean absolute error over different observations, was reached by KNN. Both bias and variances of error were proven to be reduced by KNN learning compared to model fitting. Analysis also specifies that the KNN method is more stable in the presence of decision noise, especially when all fitting based features are excluded from feature space.

We investigated the effect of learning rate (α) and Boltzmann inverse temperature (β) on model fitting error. Learning rate controls the effectiveness of new trial in comparison with the previous estimation. Low α values mean that previous estimation is precise enough for decision making, so the new observation for rewarded or unrewarded action makes slight changes in the estimation. This slight change results in the same behavior on MB and MF system. This can be attributed to the fact that differences in the environment transition probabilities, which are the most important factor that dissociate between MB and MF systems, change slowly. In addition, for too low α values, most of the time the wrong action was selected and performance of learner became non-satisfactory.

Boltzmann inverse temperature, on the other hand, controls the exploration-exploitation tradeoff. The low β values results in the similar choice probabilities for actions regardless of their values, which means more exploration. In this case, the effect of actions’ values which were calculated by either MB or MF systems decreases and are marginally (β became zero) ignored. So, it is expected that explorative subject has slight information about the preference of MB or MF system and extracting *w* will be more difficult by any estimation. High β values, show that even slightly higher values of action, make them more preferred choices which is an indication of the exploitative behavior. For higher β values, either little or huge differences in actions-values have the same effects on the observer behavior.

The proposed method is advantageous due to its lower error for extreme cases. Such extreme cases may be prevalent in clinical trials and psychiatric conditions and will make the proposed method to have superior performance over just model-fitting approaches. MAP estimation is better than MLE in extreme values due to using a prior, KNN method works very better than MAP too. The mentioned improvements will enhance the applicability of the task of Daw et. al. for computational psychiatry purposes.

We showed that using the proposed method can help to increase the statistical power in analyzing the relation between parameters such as the gaze distribution to habitual and goal directed behavior. It was proven that consideration of behavioral parameters in estimation of combination weight (in addition to fitting), improves the consistency of behavior and subjects grouping and so other conclusions from this grouping can be more precise.

It should be noted that, any model fitting tries to minimize an objective function to extract the behavior under different assumptions. The MLE maximizes the likelihood while the extracted parameter by KNN will not maximize the likelihood even if the error of extracted *w* is lower. In fact, the flow of probabilities in reinforcement agent decisions cause that a specific parameter does not guarantee maximum likelihood while another parameter exists which satisfies the maximizes likelihood criterion. Although the Cramer-Rao Lower Band can be theoretically obtained by MLE, the above statement ensures that better estimations can be obtained by learning.

The proposed method can be considered as a maximum likelihood estimation using simulation-based estimation. Such a method not only uses trial by trial observations of the behavior, but also uses global observation such as stay probabilities in random variable space and tries to maximize the likelihood of observing all the mentioned behaviors together. For large sample sizes, it is possible that MLE and KNN methods converge to the same estimation error. For limited sample sizes, however, KNN has shown more reliability and avoids overfitting, and is considered a better option in a typical experimental condition.

In sum, our proposed method can enhance the estimation of the combination weight between model based and model free behaviors. This improvement is due to using behavioral indices from the data that make the estimation more robust. This robust estimation can facilitate the using of similar paradigms in clinical applications and help in diagnosis of psychiatric disorders.

